# CBRPP: a new RNA-centric method to study RNA-protein interactions

**DOI:** 10.1101/2020.04.09.033290

**Authors:** Yunfei Li, Shengde Liu, Lili Cao, Yujie Luo, Hongqiang Du, Siji Li, Fuping You

## Abstract

RNA-protein interactions play essential roles in tuning gene expression at RNA level and modulating the function of proteins. Abnormal RNA-protein interactions lead to cell dysfunction and human diseases. Therefore, mapping networks of RNA-protein interactions is crucial for understanding cellular mechanism and pathogenesis of diseases. Different practical protein-centric methods for studying RNA-protein interactions has been reported, but few RNA-centric methods exist. Here, we developed CRISPR-based RNA proximity proteomics (CBRPP), a new RNA-centric method to identify proteins associated with the target RNA in native cellular context without cross-linking or RNA manipulation in vitro. CBRPP is based on a fusion of dCas13 and proximity-based labeling (PBL) enzyme. dCas13 can deliver PBL enzyme to the target RNA with high specificity, while PBL enzyme labels the surrounding proteins of the target RNA, which are then identified by mass spectrometry.

## Introduction

RNA is bound to protein from birth to death. RNA-binding proteins (RBPs) play a pivotal role in a wide range of biological processes, including RNA transcription, processing, modification, transport, translation and stabilization^[1–4]^. RNAs, in turn, influence proteins expression, localization and interactions with other proteins^[5–7]^. Aberrant RNA-protein interactions are related to cellular dysfunction and human diseases^[3, 8, 9]^. Therefore, mapping networks of RNA-protein interactions is of great importance for understanding many cellular biological processes.

Based on the type of molecule they start with, methods for studying RNA-protein interactions are classified into protein-centric methods and RNA-centric methods^[10]^. Protein-centric methods start with a protein of interest and study RNAs that interact with that protein. Since proteins are easily purified with antibodies, many protein-centric methods, such as cross-linking immunoprecipitation (CLIP)-seq^[11]^ and RNA immunoprecipitation (RIP)-seq^[12]^, are available and practical. Conversely, RNA-centric methods start with an RNA of interest and focus on proteins that bind it. Most approaches use biotinylated RNA^[13]^, aptamer-tagged RNA^[14]^, peptide nucleic acid^[15]^ and antisense probe^[16–20]^for purification of RNA-protein complexes to identify proteins that associate with the target RNA, however these methods often require RNA manipulation in vitro and miss transient or weak interactions. Meanwhile, compared with protein-centric methods, there are few robust RNA-centric methods.

In this paper, by combining the power of CRISPR-Cas13^[21]^ and proximity-based labeling (PBL) technique^[22]^, we developed CBRPP (CRISPR-based RNA proximity proteomics), a new RNA-centric method to identify proteins associated with the target RNA in native cellular context without cross-linking or RNA manipulation in vitro.

## Results

### Strategies to develop CBRPP

In recent years, PBL has emerged as a powerful complementary approach to classic affinity purification of multiprotein complexes in mapping of protein-protein interactions^[23]^. By fusing proteins of interest to enzymes that generate reactive molecules, most commonly biotin, adjacent proteins are covalently labeled so that they can be isolated and identified^[22]^. To date, multiple versions of the PBL enzyme have been developed, such as BioID2^[24]^, TurboID^[25]^, Apex2^[26]^ and BASU^[27]^. The key advantage of PBL is that it can capture weak and transient interactions in live cells. Recently, two studies have applied PBL to study RNA-protein interactions using the MS2-MCP strategy^[28]^ or a similar strategy^[27]^, but both require insertion of MS2 or BoxB stem-loop into the target RNA in advance, which may influence structure or function of the target RNA.

The discovery of RNA-targeting CRISPR systems offers scientists a powerful toolbox to manipulate RNA in live cells^[21]^. Active Cas13, under the guidance of the specific CRISPR RNA (crRNA), can recognize and cleave the target RNA. Catalytically dead Cas13 (dCas13) retains programmable RNA-binding capability, which can be utilized for RNA imaging and editing^[29, 30]^. Currently, there are several orthologs and subtypes of Cas13 that are catalytically active inside mammalian cells, including LwaCas13a^[29]^, PspCas13b^[30]^ and RfxCas13d^[31]^. Inspired by GLoPro^[32]^ and C-BERST^[33]^, we proposed that by fusing dCas13 and PBL enzyme together, dCas13, under the guidance of a specific crRNA, can act as an RNA tracker to bring PBL enzyme to the target RNA, then PBL enzyme can biotinylate the surrounding proteins of the target RNA with biotin. Finally, these biotinylated proteins can be easily enriched by streptavidin beads and identified by liquid chromatography mass spectrometry (LC-MS) (Figure 1). We referred to this combination of CRISPR-Cas13 and PBL as CRISPR-based RNA proximity proteomics (CBRPP).

**Figure 1.**
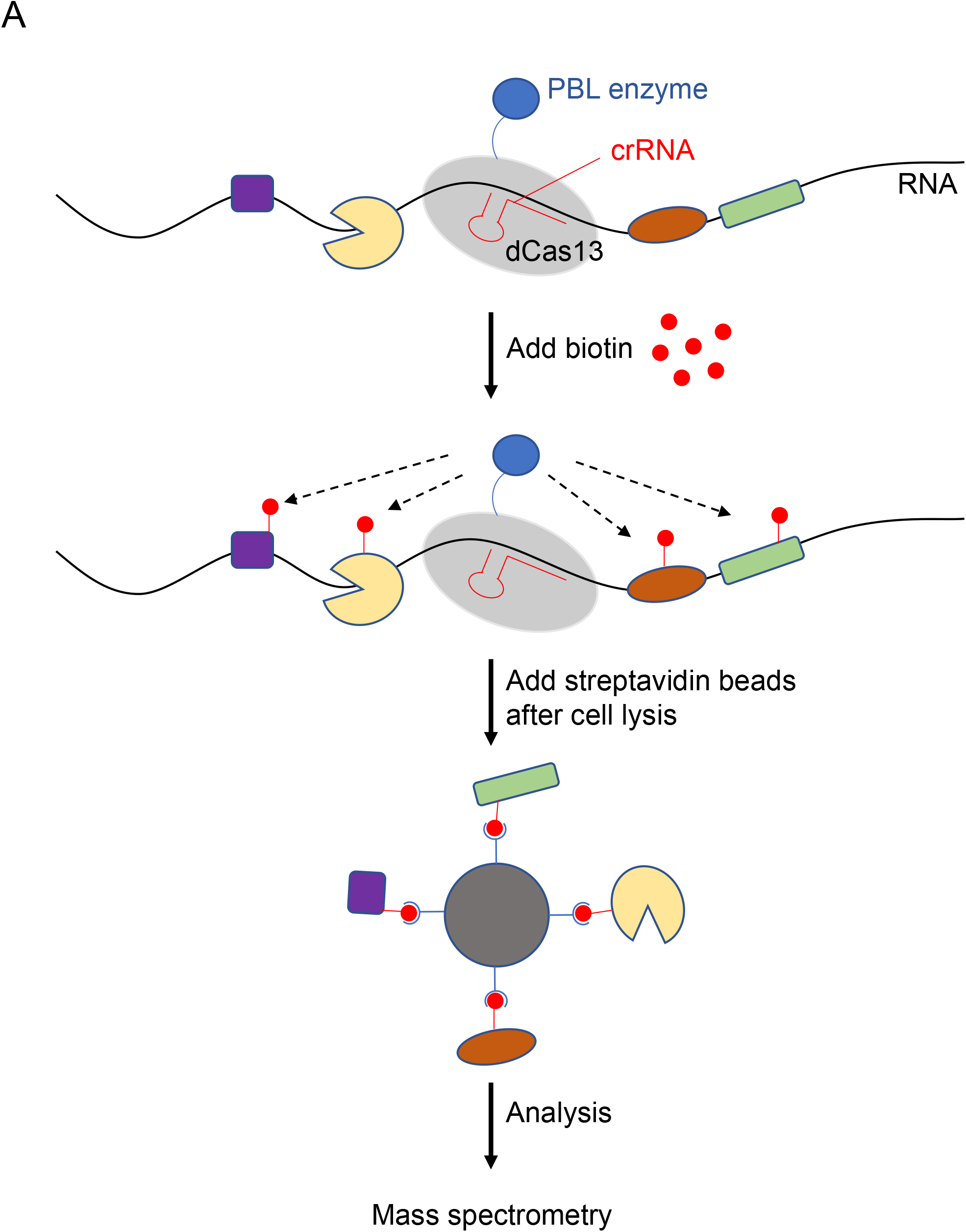
Design of CBRPP. (A) Schematic representation of CBRPP approach. By fusing dCas13 and PBL enzyme together, dCas13, under the guidance of a specific crRNA, acts as an RNA tracker to bring PBL enzyme to the target RNA, then PBL enzyme biotinylates surrounding proteins of the target RNA, followed by streptavidin beads enrichment of biotinylated proteins and mass spectrometry.

### dRfxCas13d is not suitable for CBRPP to study RNA-protein interactions

To prove the concept, we firstly selected dRfxCas13d and APEX2 for testing, because RfxCas13d is the smallest and most active one among Cas13 proteins^[31]^ and APEX2 have the fastest rate of labeling^[26]^, which can be used for isolated analysis of RNA-protein interactions that occur over short time periods. We fused APEX2 to N-terminus or C-terminus of RfxCas13d to test whether the fusion of APEX2 affected the function of RfxCas13d by detecting knockdown efficiency of RfxCas13d (Figure 2A). Results showed that fusion of APEX2 to C-terminus of RfxCas13d only slightly affect the knockdown efficiency of RfxCas13d, and has no effect on the expression of RfxCas13d (Figure 2B). Therefore, we constructed dRfxCas13d-APEX2-NES plasmid (Figure 2A) and applied it to well-studied ACTB mRNA to test whether it would identify known RBPs of ACTB mRNA. We designed seven Rfx-crRNAs targeting different regions of ACTB mRNA and validated their targeting by knockdown with an active RfxCas13d. RT-qPCR results showed that all seven Rfx-crRNAs significantly reduced ACTB mRNA levels in HEK293T cells (Figure 2C). Then we transfected HEK293T with dRfxCas13d-APEX2-NES and two optimal ACTB Rfx-crRNAs (crRNA4 and crRNA7) to test whether dRfxCas13d-APEX2-NES can be directed to ACTB mRNA under the guidance of ACTB Rfx-crRNAs. In addition, cells were treated with sodium arsenite to induce the formation of stress granules where ACTB mRNA accumulated. Results showed that dRfxCas13d-APEX2-NES could colocalize with the stress granule marker G3BP1 regardless of co-transfection with ACTB Rfx-crRNAs or non-targeting Rfx-crRNAs (Figure 2D). This indicated that dRfxCas13d-APEX2-NES may nonspecifically accumulate with ACTB mRNA. We also constructed dRfxCas13d-APEX2-NLS plasmid and designed the Rfx-crRNAs targeting NEAT1 to study paraspeckles. We found that the localization of dRfxCas13d-APEX2-NLS had no difference between the non-targeting crRNA group and the NEAT1 targeting crRNA group (data not show), which is consistent with results of Chen lab^[34]^. These data suggested that dRfxCas13d is not suitable for CBRPP to study RNA-protein interactions.

**Figure 2.**
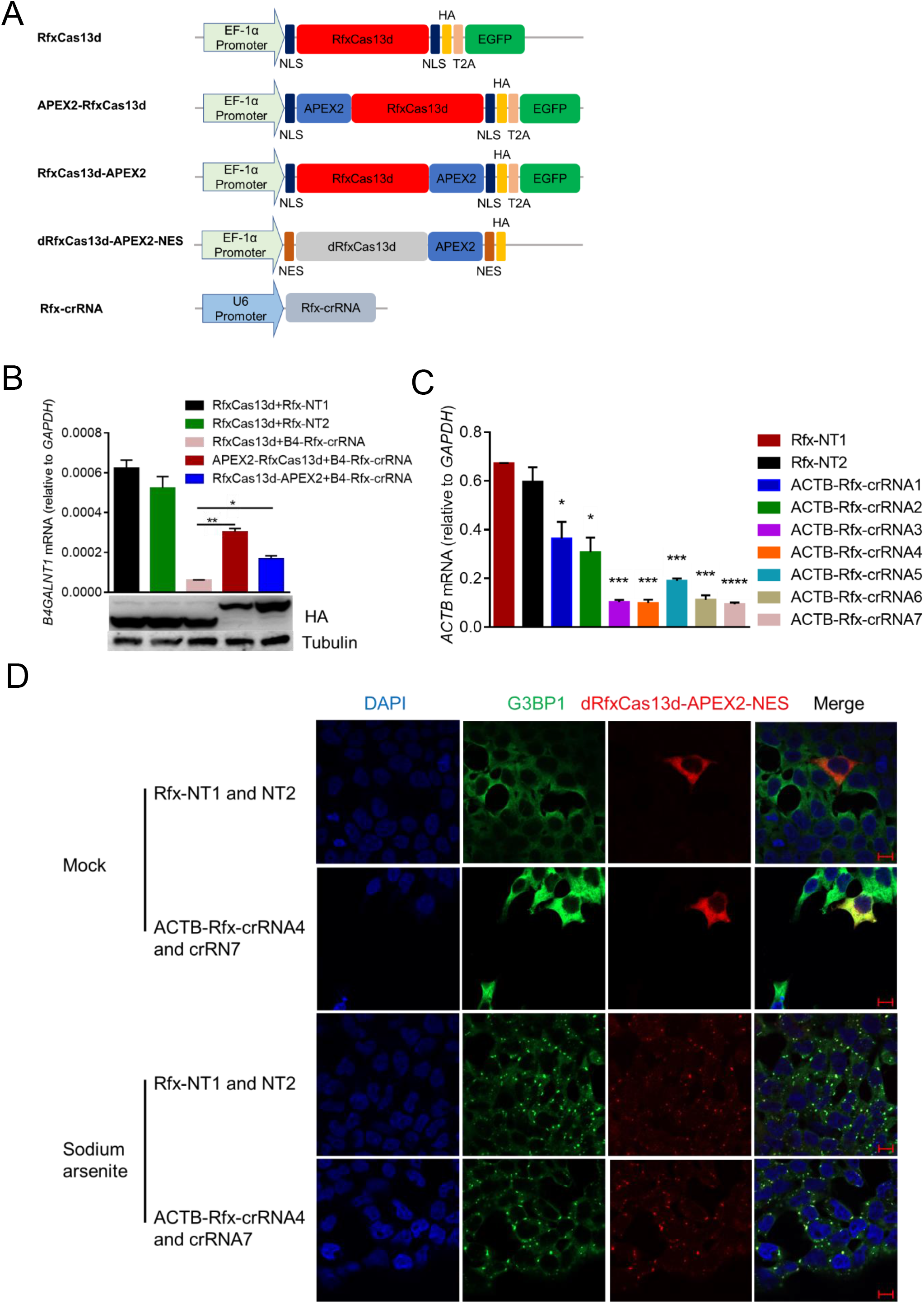
dRfxCas13d is not suitable for CBRPP to study RNA-protein interactions. (A) Plasmids used in this figure. NLS: nuclear localization sequence; NES: nuclear export sequence; EGFP: enhanced green fluorescent protein; T2A: T2A self-cleaving peptide; HA: hemagglutinin tag. (B) Upper: HEK293T cells were co-transfected with RfxCas13d/RfxCas13d-APEX2/APEX2-RfxCas13d and B4GALNT1 Rfx-crRNA to detect the mRNA level of B4GALNT1 by RT-qPCR after 48 hours. Rfx-NT1: non-targeting Rfx-crRNA 1; Rfx-NT2: non-targeting Rfx-crRNA 2. B4-Rfx-crRNA: B4GALNT1 Rfx-crRNA. Bottom: western blotting to measure the protein expression level of RfxCas13d, APEX2-RfxCas13d and RfxCas13d-APEX2. (C) HEK293T cells were co-transfected with RfxCas13d and ACTB Rfx-crRNAs to detect the mRNA level of ACTB by RT-qPCR after 48 hours. (D) Representative images for dRfxCas13d-APEX2 imaging with two crRNAs targeting ACTB mRNA in HEK293T. Mock: no treatment. Sodium arsenite: treating cells with 0.5mM sodium arsenite for 30 minutes. Stress granules are indicated by G3BP1 staining. Scale bars, 10 μm.

### Transient transfection of dPspCas13b-APEX2 to study RBPs of ACTB mRNA is not effective

Recent study showed that dPspCas13b is the most efficient dCas13 protein to label RNA^[34]^, so we replaced dRfxCas13d with dPspCas13b and added a flexible linker 3x(GGGGS) between dPspCas13b and APEX2 to avoid mutual influence (Figure 3A). Since PspCas13b and RfxCas13d cannot share the crRNAs, we redesigned four ACTB Psp-crRNAs and validated their targeting. Results showed that all four ACTB Psp-crRNAs significantly reduced ACTB mRNA levels in HEK293T cells, and that the knockdown efficiency and expression level were comparable between PspCas13b and PspCas13b-APEX2 (Figure 3B and 3C). Furthermore, co-transfection of dPspCas13b-APEX2 and ACTB Psp-crRNAs in HEK293T did not affect the mRNA and protein level of ACTB (Figure 3D), suggesting this system does not affect the stability and translation of ACTB mRNA. Then, we transiently transfected dPspCas13b-APEX2 and ACTB Psp-crRNAs into HEK293T cells to performed a 1-minute proximity labeling reaction, followed by streptavidin bead enrichment of biotinylated proteins and LC-MS (Figure 3E). The streptavidin-HRP blot showed that dPspCas13b-APEX2 has biotinylation activity (Figure 3F). Mass spectrometry profiling results showed that the known RBPs of ACTB mRNA (marked in red) were not enriched in the ACTB Psp-crRNA group relative to the non-targeting Psp-crRNA group (Figure 3G). These data suggested that transient transfection of dPspCas13b-APEX2 to study RBPs of ACTB mRNA is not effective.

**Figure 3.**
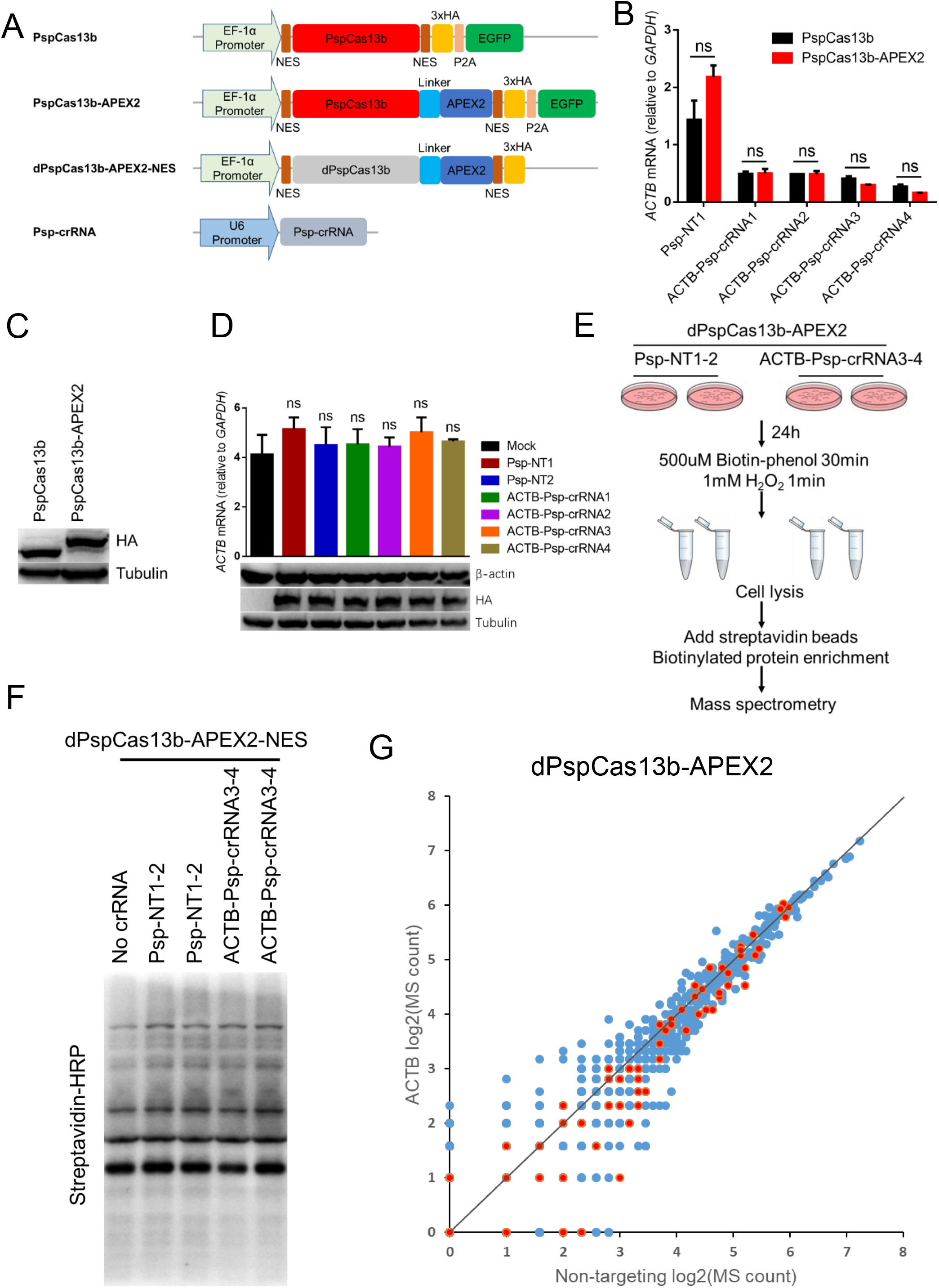
Transient transfection of dPspCas13b-APEX2 to identify RBPs of ACTB mRNA. (A) Plasmids used in this figure. P2A: T2A self-cleaving peptide; Linker: 3x(GGGGS), G: glycine, S: serine. (B) HEK293T cells were co-transfected with PspCas13b/PspCas13b-APEX2 and ACTB Psp-crRNAs to detect the mRNA level of ACTB after 48 hours. Psp-NT1: non-targeting Psp-crRNA 1; Psp-NT2: non-targeting Psp-crRNA 2. (C) Western blotting to measure the protein expression level of PspCas13b and PspCas13b-APEX2. (D) HEK293T cells were co-transfected with dPspCas13b-APEX2-NES and different ACTB Psp-crRNAs. Upper: RT-qPCR analysis of ACTB mRNA level in cells. Bottom: western blotting to measure the protein expression level of ACTB in cells. (E) Workflow of transiently transfected dPspCas13b-APEX2 to capture the proteins that interact with ACTB mRNA in HEK293T. Psp-NT1-2: non-targeting Psp-crRNA 1 and non-targeting Psp-crRNA 2; ACTB-Psp-crRNA3-4: ACTB Psp-crRNA 3 and ACTB Psp-crRNA 4. (F) Western blotting to detect the biotinylation activity of HEK293T cells co-transfected with dPspCas13b-APEX2-NES and different Psp-crRNAs. (G) Scatter plot showing the number of peptides per protein after log2 transformation in non-targeting Psp-crRNA group (X-axis) and ACTB Psp-crRNA group (Y-axis) from mass spectrometry proteomics data. The red dots in the scatter plot represent known RBPs of ACTB mRNA in StarBase v2.0 database. The experiments were done in HEK293T transiently transfected with dPspCas13b-APEX2 and Psp-crRNAs.

We speculated that such results may be due to the high expression of dPspCas13b-APEX2 or the properties of APEX2 itself. If the protein expression level of dPspCas13b-APEX2 is too high, or the copy number of dPspCas13b-APEX2 proteins exceeds that of target RNAs, some redundant dPspCas13b-APEX2 proteins cannot be directed to the target RNAs with the guidance of specific crRNAs, so background proteins would be labeled, resulting in low signal-to-noise ratio. It’s known that APEX2-based labeling is often specific to low-abundance amino acids such as tyrosine^[35, 36]^, so it is possible that labeling will not occur if surface-exposed tyrosine is not available.

### Inducible expression of dPspCas13b-BioID2 successfully identifies RBPs of ACTB mRNA

For further optimization, we next used other three PBL enzymes (BioID2, TurboID and BASU) to test which enzyme is optimal (Figure 4A). Simultaneously, we took advantage of the Tet-On 3G inducible expression system to keep the expression of fusion proteins at a low level in HEK293T cells (Supplementary Figure 1). RT-qPCR results showed that the knockdown efficiency of PspCas13b was not affected by fusion BioID2/TurboID/BASU/APEX2 to C terminus of PspCas13b (Figure 4B). Therefore, we constructed four stable HEK293T cell lines for inducible expression of dPspCas13b-BioID2/TurboID/BASU/Apex2. Western blotting results showed that all four stable cell lines can be induced by doxycycline in a dose-dependent manner, and that BioID2, TurboID and APEX2 have biotinylation activity but not BASU (Figure 4C). Subsequently, we used dPspCas13b-BioID2/TurboID/Apex2 inducibly expressing cell lines to identify the proteins interacting with ACTB mRNA (Figure 4D and 4E). We analyzed the protein mass spectrometry data obtained from these three cell lines, and found that in dPspCas13b-BioID2 inducibly expressing cell line, the known RBPs of ACTB mRNA (marked in red) such as IGF2BP1, HNRNPA1, HNRNRC, HNRNPA2B1 and HNRNPM, were significantly enriched in ACTB Psp-crRNA group relative to the non-targeting Psp-crRNA group (Figure 4F and Supplementary Figure 2). IGF2BP1, also known as ZBP1 (zipcode-binding protein 1), interacts with the zipcode of ACTB mRNA via KH (HNRNPK homology) domains to regulate the localization and translation of ACTB mRNA^[37]^. HNRNPA1, HNRNPC, HNRNPA2B1 and HNRNPM are common RBPs that are involved not only in processing heterogeneous nuclear RNAs (hnRNAs) into mRNAs, but also mRNAs stability and translational regulation^[38]^. KHSRP have been suggested to be associated with ACTB mRNA localization^[39]^. These data indicated that inducible expression of dPspCas13b-BioID2 successfully identify RBPs of ACTB mRNA.

**Figure 4.**
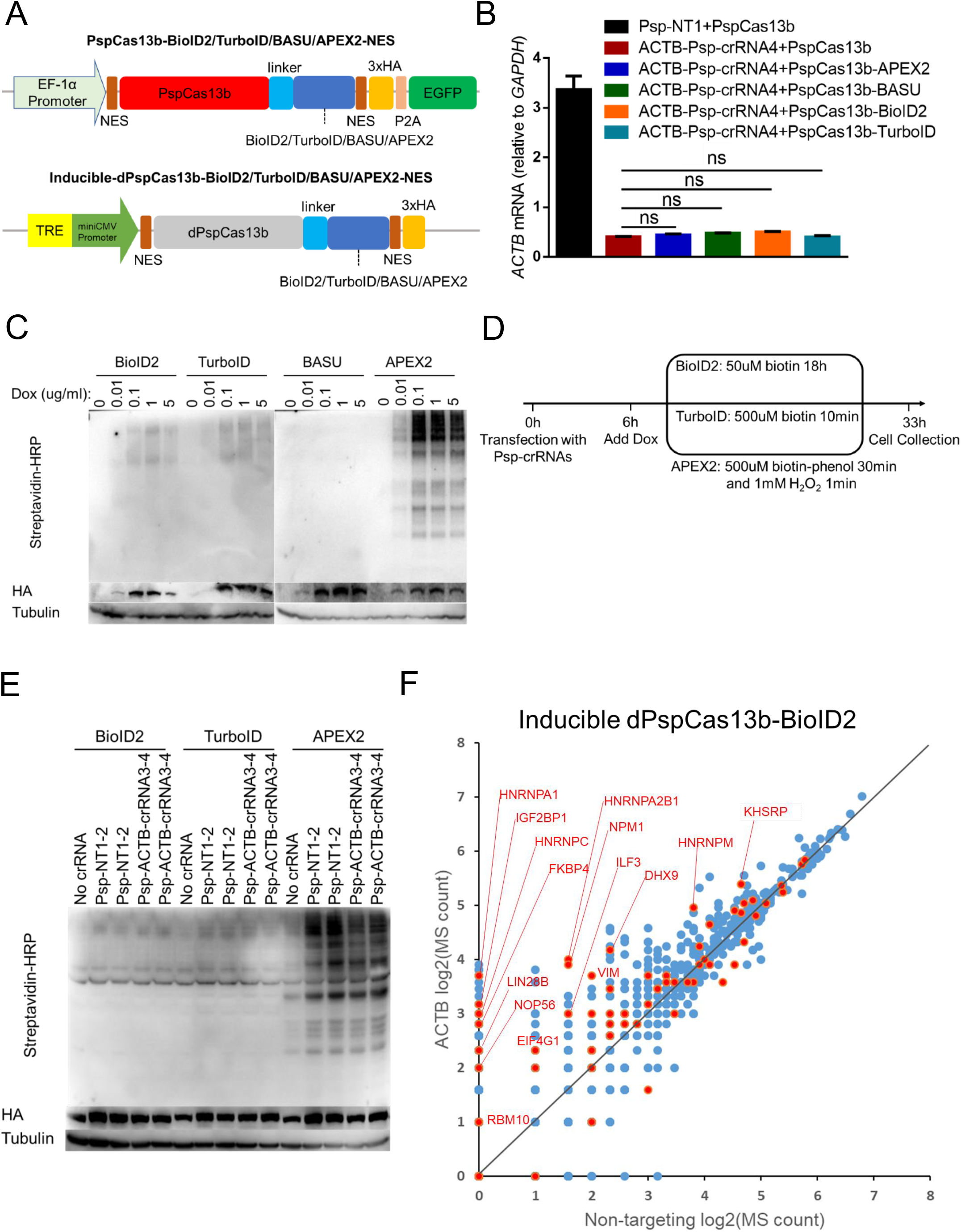
Using dPspCas13b-BioID2/TurboID/APEX2 inducibly expressing cell lines to identify RBPs of ACTB mRNA. (A) Plasmids used in this figure. (B) HEK293T cells were co-transfected with PspCas13b or PspCas13b-APEX2/BASU/BioID2/TurboID and ACTB Psp-crRNAs to detect the mRNA level of ACTB after 48 hours. (C) Western blotting to test the inducible ability and the biotinylation activity of four stable HEK293T cell lines for inducible expression of dPspCas13b-BioID2/TurboID/BASU/Apex2. Dox: doxycycline. (D) Timeline to capture the proteins that interact with ACTB mRNA using dPspCas13b-BioID2/TurboID/Apex2 inducibly expressing cell lines. (E) Western blotting to detect the expression level and biotinylation activity of cells collected from (D). (F) Scatter plot showing the number of peptides per protein after log2 transformation in non-targeting Psp-crRNA group (X-axis) and ACTB Psp-crRNA group (Y-axis) from mass spectrometry proteomics data. The red dots in the scatter plot represent known RBPs of ACTB mRNA in StarBase v2.0 database. The experiments were done in dPspCas13b-BioID2 inducibly expressing cell line.

Unlike TurboID or APEX2, BioID2 used in CBRPP generates a history of RNA-protein interactions over time, which can capture some transient RNA-protein interactions, such as those occur during various stages of the cell cycle. Besides, the results obtained using BioID2 in CBRPP represent the accumulation of biotinylated proteins over the labeling time. The proteins that interact with the target RNA are labeled and accumulated during this time, and those background proteins that occasionally appear near the target RNA without mutual interaction may be labeled but not accumulated, which result in high signal-to-noise ratio. Therefore, inducible expression of dPspCas13b-BioID2 is recommended to study RBPs of the target RNA.

## Discussion

Here we proposed a new RNA-centric method named CBRPP by combining dCas13 with proximity-based labeling. With some optimizations, we finally determined that inducible expression of dPspCas13b-BioID2 is most suitable for studying RNA-protein interactions. In the presence of a specific crRNA, the dPspCas13b-BioID2 fusion protein is directed to the target RNA, then BioID2 in the chimera biotinylates nearby proteins of the target RNA. With the strong interaction between biotin and streptavidin, biotinylated proteins can be easily enriched and identified.

Compared with previous RNA-centric methods, CBRPP has several advantages. First, CBRPP does not require pre-labeling of the target RNA^[13]^, MS2 insertion in advance^[28]^, or designing antisense probes^[16–20]^ to purify RNA-protein complexes. In dPspCas13b-BioID2 positive cells only crRNAs are required. Second, using CBRPP, RBPs labeling is done in a living cell state without manipulating RNA-protein complexes in vitro, so it almost preserves the natural structure of the target RNA, while avoiding the possible disruption of RNA-protein interactions and RNA degradation. Third, CBRPP can capture weak and transient RNA-protein interactions, taking advantage of proximity-based labeling^[22]^.

As with any technology, CBRPP has its limitations. Since proximity-based labeling is in a distance-dependent manner, proteins identified by CBRPP may be not RBPs of the target RNA but merely proximate proteins. Therefore, it is necessary to confirm the interactions between the target RNA and candidate proteins identified by CBRPP with RIP or CLIP. Due to the large size of dPspCas13b-BioID2, its binding to the target RNA may affect the binding of the original interacted protein at this site. In addition, the long labeling time required for BioID2 methods prevents CBRPP from isolated analysis of RNA-protein interactions that occur over short period of time.

According to our experience, there are three crucial factors for the success of CBRPP. First, it is necessary to find potent crRNAs for analysis, and testing multiple crRNAs at the same time is recommended. Second, the expression level of dPspCas13b-BioID2 should be controlled at a low level in case the copy number of fusion proteins exceeds that of the target RNAs, resulting in low signal-to-noise ratio. Third, setting up an appropriate control group is very helpful for excluding background proteins identified by the experimental group.

In summary, in this study we developed an effective RNA centric method to identify proteins associated with the target RNA in native cellular context without cross-linking or RNA manipulation in vitro. Although we have only studied ACTB mRNA using CBRPP, in principle CBRPP can also be used to study lncRNA or other RNA types. For large lncRNA, taking Xist as an example, by designing different crRNAs target different regions of Xist, CBRPP can not only study the RBPs of Xist, but also the RBPs at a certain position of Xist^[16, 40]^. Furthermore, CBRPP is suitable for studying the mechanism of diseases caused by abnormal RNA, such as myotonic dystrophy type 1^[7]^.

## Materials and Methods

### Cell culture

HEK293T (Human Embryonic Kidney 293T) cells was obtained from ATCC. Cells were cultured in DMEM medium supplemented with 10% FBS (Gibco) and 100U/ml Penicillin-Streptomycin in a humidified incubator at 37 °C with 5% CO_2_.

### Reagents and Antibodies

PEI (764582, Sigma-Aldrich) was used for transfection. Antibodies used in this study include the following: anti-HA (rabbit, H6908, Sigma-Aldrich); anti-alpha-tubulin (rabbit, 11224-1-AP, Proteintech); anti-G3BP1 (mouse, sc-365338, Santa Cruz); HRP-conjugated Streptavidin (SA00001-0, Proteintech). The antibodies were diluted 1,000 times for immunoblots, 200 times in confocal microscopy.

### Plasmid constructs

Expression constructs generated for this study were prepared by standard molecular biology techniques and coding sequences entirely verified. All the mutants were constructed by standard molecular biology technique. Each mutant was confirmed by sequencing. All plasmid constructs and their sequence were listed in Supplementary Table 1. All crRNAs used in this paper were listed in Supplementary Table 2.

### Western blotting

Cells were washed with PBS and lysed by incubation on ice for 10 min with RIPA lysis buffer (50 mM Tris, 150 mM NaCl, 0.1% SDS, 0.5% sodium deoxycholate, 1% Triton X-100, protease cocktail [C0001, Targetmol], and 1 mM PMSF). The proteins were resolved by SDS/PAGE and transferred to 0.22 um nitrocellulose membrane (PALL), which then was incubated overnight with primary antibodies. The membrane was further incubated with the corresponding HRP-conjugated secondary antibodies and detected by enhanced chemiluminescence.

### Immunofluorescence microscopy

HEK293T cells were plated and grew on coverslips with indicated treatments, washed with pre-warmed PBS, and fixed with 4% paraformaldehyde for 10 min. The cells were permeated with 0.5% Triton-100 for 3 min, blocked with 3% BSA for 30 min, washed, and incubated with primary antibodies for 1 h at 37 °C. After washing, cells were stained with Alexa Fluor 488-conjugated secondary antibodies (A11029, Invitrogen) or Alexa Fluor 555-conjugated secondary antibodies (A-21428, Invitrogen) for 1 h at 37 °C, and then with DAPI (4′,6-Diamidino-2-phenylindole, Roche) for 15 min. The coverslips were washed extensively and mounted onto slides. Imaging of the cells were carried out using N-STORM5.0 microscope.

### RNA Extraction and Quantitative reverse transcription PCR (RT-qPCR)

Total RNA from cells were isolated using the RNA simple Total RNA kit (TIANGEN). 1ug RNA was reverse transcribed using a FastKing RT Kit (TIANGEN). Levels of the indicated genes were analyzed by quantitative real-time PCR amplified using SYBR Green (Q311, Vazyme). All primers were listed in Supplementary Table 3.

### Generation of Stable Expression Mammalian Cell Lines

For preparation of lentiviruses, HEK293T cells in 6-well plates were transfected with the lentiviral vector of interest (1,800 ng), the lentiviral packaging plasmids psPAX2 (600 ng) and pMD2.G (600 ng) and 12 ul of PEI (1mg/ml). About 48 h after transfection the cell medium containing lentiviruses was centrifugalized at 12,000 g for 3 minutes and the supernatant was harvested. HEK293 cells were then infected at ~50% confluency by lentiviruses for 48 h, followed by selection with 1 μg/ml puromycin in growth medium for 7 days. The stable transgene monoclonal cells were harvested by limiting dilution in cell pools.

### Generation of Tetracycline (Tet) Inducible Expression HEK293T cell lines

The two consecutive manipulation steps are necessary to generate human Tet-on cell lines with inducible expression of plasmids of interest. The first step is generation of cells stably expressing reverse tetracycline-controlled transactivator (rtTA). HEK293T cells were infected at ~50% confluency by lentiviruses containing pLVX-TetO3G(rtTA)-hygr vector for 48 h, followed by selection with 50 ug/ml hygromycin in growth medium for 7 days, and hygromycin resistant clones were selected. Several clones were picked and tested for rtTA expression by immunoblotting. After testing for all molecular and cell biological parameters of interest, the ‘best’ rtTA-positive clone was expanded and stored. The next step is generation of Tet-on cell lines with inducible expression of target plasmids. The ‘best’ rtTA-positive clone was infected by lentiviruses containing target plasmids (Inducible-dPspCas13b-BioID2/BASU/TurboID/APEX2) for 48 h, followed by selection with 1 μg/ml puromycin in growth medium for 7 days. The puromycin resistant clones were harvested by limiting dilution in cell pools. Several individual cell clones were picked, expanded and screened by immunoblotting for Doxycycline-inducible expression of the gene of interest. Finally, clones of interest were expanded, re-tested and stored.

### Biotin Labeling in Live Cells

For dPspCas13b-APEX2 transient transfection experiments, HEK293T cells were plated in 10 cm dish at 70% confluency 18 hours prior to transfection. Cells were transfected with the dPspCas13b-Apex2 plasmid and the crRNA plasmids. After 6 hours of transfection, the culture medium was changed. After 24 hours of transfection, biotin-phenol was added to cell culture medium to a final concentration of 500 uM for 30 minutes, H_2_O_2_ was then added into cell culture media at a final concentration of 1mM to induce biotinylation. After gently shaking the cell culture dish for one minute, the medium was removed and cells were washed three times with PBS supplemented with 100mM sodium azide, 100mM sodium ascorbate and 50mM TROLOX. Cells were scraped and transferred to 1.5 ml tubes with ice cold PBS, spun at 3600 rpm for 5 minutes, flash frozen in liquid nitrogen and stored at −80°C.

For inducibly expressing dPspCas13b-BioID2/TurboID/BASU/Apex2 experiments, four stable HEK293T cell lines for inducible expression of dPspCas13b-BioID2/TurboID/BASU/Apex2 were plated in 10 cm dish at 70% confluency 18 hours prior to transfection. Cells were transfected with 20ug crRNA plasmid per dish. After 6 hours of transfection, the culture medium was replaced with new media containing 0.1 ug/ml doxycycline. For BioID2, biotin was added to the culture medium at a final concentration of 50 uM after 15 hours of transfection; for TurboID, biotin was added at a final concentration of 500uM for 10 minutes before harvesting cells; for BASU, biotin was added at a final concentration of 200uM for 2 hours before harvesting cells; for APEX2, biotin-phenol was added at a final concentration of 500 uM for 30 minutes and H_2_O_2_ was added at a final concentration of 1 mM for one minute before harvesting cells. All kinds of cells were harvested at 33 hours after transfection. For APEX2, the medium was removed and cells were washed three times with ice cold PBS supplemented with 100 mM sodium azide, 100mM sodium ascorbate and 50mM TROLOX; for TurboID/BASU/BioID2, the medium was removed and cells were washed three times with ice cold PBS. Cells were scraped and transferred to 1.5 ml tubes with ice cold PBS, spun at 3600 rpm for 5 minutes, flash frozen in liquid nitrogen and stored at −80°C.

### Streptavidin Magnetic Bead Enrichment of Biotinylated Proteins

Cell pellets as described above were lysed in RIPA lysis buffer (50 mM Tris, 150 mM NaCl, 0.1% SDS, 0.5% sodium deoxycholate, 1% Triton X-100, protease cocktail [TargetMol], and 1 mM PMSF) at 4°C for 10 minutes. The lysates were cleared by centrifugation at 12,000 g for 10 min at 4 °C. 50ul of each lysate supernatant was reserved for detection of biotinylation activity by western blotting. Streptavidin magnetic beads were washed twice with RIPA lysis buffer and then mixed with lysates supernatant together with rotation overnight at 4 °C. On day 2, the beads were subsequently washed twice with 1 mL of RIPA lysis buffer, once with 1 mL of 1 M KCl, once with 1 mL of 0.1 M Na_2_CO_3_, once with 1 mL of 2 M urea in 10 mM Tris-HCl (pH 8.0), and twice with 1 mL RIPA lysis buffer. Finally, biotinylated proteins were eluted by boiling the beads in 150 μL of elution buffer (55 mM pH 8.0 Tris-HCl, 0.1% SDS, 6.66mM DTT, 0.66 mM biotin) for 10 minutes and sent for mass spectrometry.

### Statistical Analysis

The descriptive statistical analysis was performed with Prism version 7 (GraphPad Software). All data are presented as mean ± SD. A two-tailed Student’s t test assuming equal variants was used to compare two groups. In all figures, the statistical significance between the indicated samples and control is designated as *P < 0.05, **P < 0.01, ***P < 0.001, or NS (P > 0.05).

## Acknowledgments

This work was supported by the National Natural Science Foundation of China (31570891) and the National Key Research and Development Program of China (Grant #2016YFA0500302).

## Author Contributions

Y.L. and F.Y. conceived this project. Y.L. analyzed the data and wrote the paper. Y.L., SD.L., L.C., H.D. and F.Y. revised the paper. Y.L., SD.L., L.C. and YJ.L. performed most experiments. SJ.L. contributed to imaging.

## Table legends

**Table 1. plasmids used in this paper**

**Table 2. crRNAs used in this paper**

**Table 3. qPCR primers used in this paper**

## Supplementary figure legends

**Supplementary Figure 1**

(A) Work model of Tet-On 3G inducible expression system. Doxycycline binds the rtTA transcription factor and allows it to bind DNA at the promoter. Gene expression is induced in the presence of doxycycline. Reverse tetracycline-controlled transactivator (rtTA) is created by fusing reverse Tet repressor (rTetR) with VP16. TRE: Tet response element. Dox: doxycycline, a analog of tetracycline.

**Supplementary Figure 2**

Scatter plot showing the number of peptides per protein after log2 transformation in non-targeting Psp-crRNA group (X-axis) and ACTB Psp-crRNA group (Y-axis) from mass spectrometry proteomics data. The red dots in the scatter plot represent known RBPs of ACTB mRNA in StarBase v2.0 database.

(A) The experiments were done in dPspCas13b-APEX2 inducibly expressing cell line.

(B) The experiments were done in dPspCas13b-TurboID inducibly expressing cell line.

