## Supplementary figures for "CBRPP: a new RNA-centric method to study RNA-protein interactions"

A

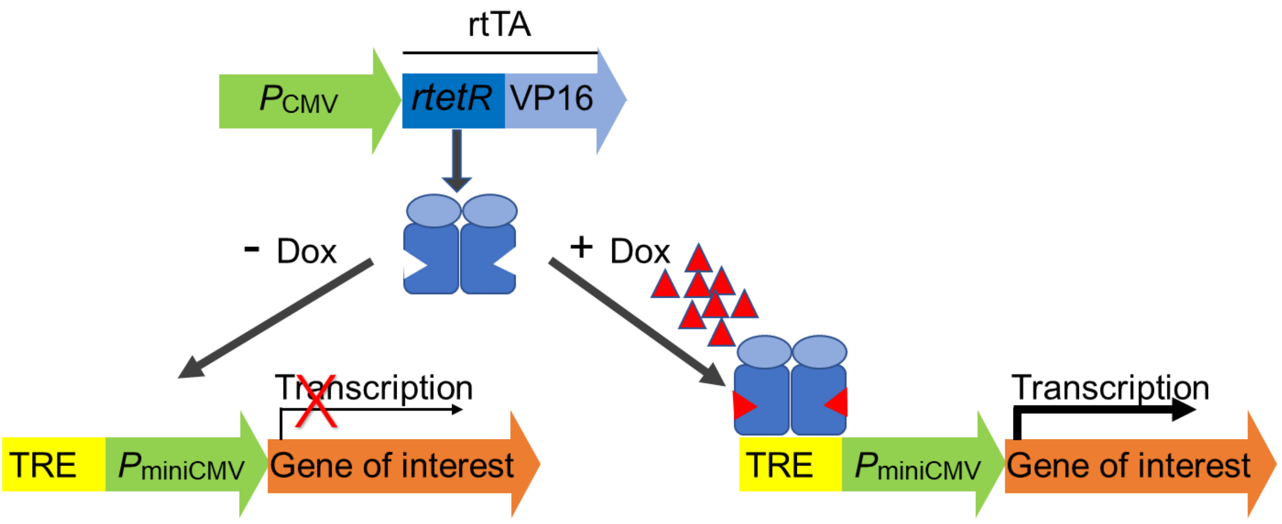


Supplementary Figure 1

A

Inducible dPspCas13b-APEX2

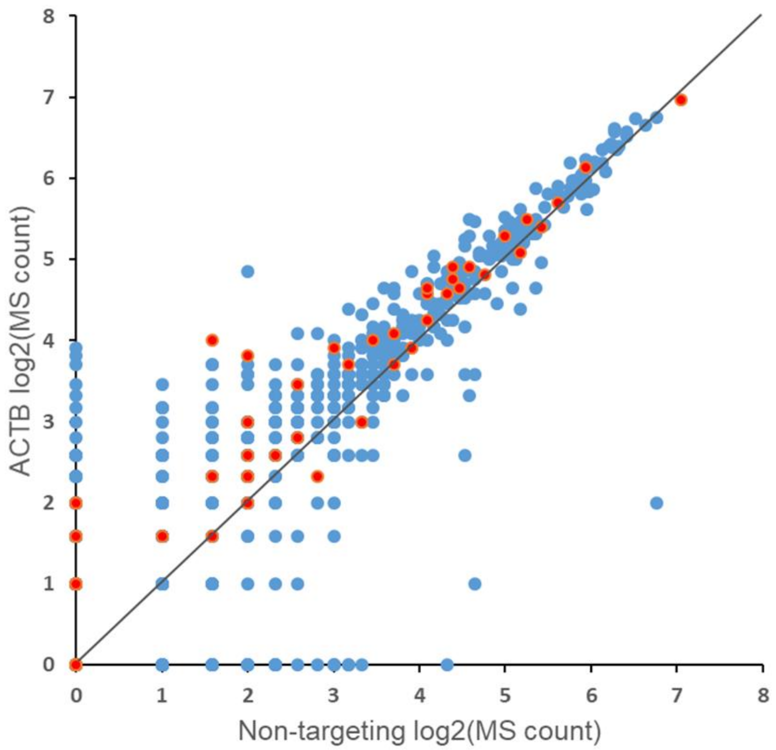


B

Inducible dPspCas13b-TurboID

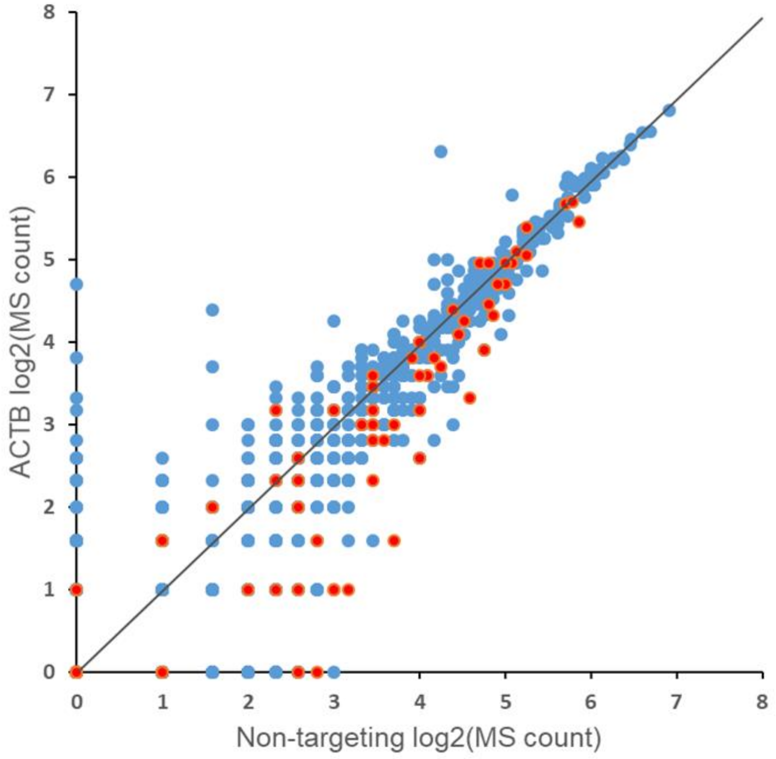


Supplementary Figure 2
